# Sequence based prediction of novel domains in the cellulosome of *Ruminiclostridium thermocellum*

**DOI:** 10.1101/066357

**Authors:** Zarrin Basharat, Azra Yasmin

**Affiliations:** Microbiology & Biotechnology Research Lab, Department of Environmental Sciences, Fatima Jinnah Women University, 46000, Pakistan

**Keywords:** Domain prediction, *in silico*, *R. thermocellum*, cellulosome

## Abstract

*Ruminiclostridium thermocellum* strain ATCC 27405 is valuable with reference to the next generation biofuel production being a degrader of crystalline cellulose. The completion of its genome sequence has revealed that this organism carries 3,376 genes with more than hundred genes encoding for enzymes involved in cellulysis. Novel protein domain discovery in the cellulose degrading enzyme complex of this strain has been attempted to understand this organism at molecular level. Streamlined automated methods were employed to generate possibly unreported or new domains. A set of 12 novel Pfam-B domains was developed after detailed analysis. This finding will enhance our understanding of this bacterium and its molecular processes involved in the degradation of cellulose. This approach of *in silico* analysis prior to experimentation facilitates in lab study. Previously uncorrelated data has been utilized for rapid generation of new biological information in this study.

**Note:** This research was conducted in 2014 for *Clostridium thermocellum* ATCC 27405. The bacterium was later reannotated as *Ruminiclostridium thermocellum*. See NCBI nonredundant RefSeq protein annotation details at http://www.ncbi.nlm.nih.gov/refseq/about/prokaryotes/reannotation/. The study utilizes Pfam-B database, which was discontinued with effect from release 28.0 (5/2015). Availability of new information, reannotation/modification in accession numbers might impact some of the analyzed values although effort has been made to provide latest accession numbers and reference strain parameters. The preprint version may contain grammatical and proofreading mistakes. Errors and omissions excepted.

## Introduction

*Ruminiclostridium thermocellum* ATCC 27405 is a gram positive, rod shaped thermophilic bacteria which is anaerobic and produces spores [1]. The genome of this bacterium was completely sequenced by DOE Joint Genome Institute as it contains a unique extracellular enzyme system (cellulosome) gifted for breaking down insoluble cellulose into ethanol which is crucial for biomass energy production[2]. The cellulosome of this bacterial strain consists of almost 20 catalytic enzymes encoded by more than 100 genes. Some genes exercise synthesis of the enzymes while others regulate cellulosome activity with the help of inducers and repressors. Enhancement and engineering of these enzymes can lead to more efficient degradation and production of soluble sugars, ethanol and other useful chemicals and hence play a role in the biofuel production. Also, due to the potential of *R. thermocellum* to degrade cellulose into fermentative products, it has an effect on the environment by contributing to the carbon cycle and decomposition of biomass [3].

Understanding of the molecular biology of an organism by comparison of the proteomes is a useful tool in finding close and direct relations but it also misses the subtler associations between proteins. An extra refined method of analyzing proteins is through the determination of their domain content [4]. Protein domains are discrete stable amino acid structures, usually globular and created between 40 and 400 amino acids. Homologous domains reveal extremely alike tertiary structures, with the overall structure of the protein being a combination of its domains and connecting segments. Biochemical and physiological functions may also be conveyed between homologous domains to a varying extent. A wide-range of interactions, specificities or activities are exhibited by some domain families, whereas others are confirmed to demonstrate far less variation. Analogous to domains are structural repeats, normally between 5 and 60 amino acid residues in length. Repeats exist as a tandem array in the protein and fold together to form stable three dimensional structures [5]. Repeats are different from repeated domains. It is anticipated that repeated domains are stable in isolation, contrasted to the repeats which are not [6].

The proteome of *R. thermocellum* consists of a set of 3,224 proteins with 1,763 conserved domains. The proteins of our interest are enzymes which play an important role in cellulysis and maybe further studied to engineer the strain for enhanced cellulose degradation for biofuel production. We were tempted to use the kitchen sink approach to narrow down the number of databases to be used in the domain scanning process. The scanning of sequence against the large number of databases only slows down the search process and gives repeated results. In case of any doubt, the subset of sequence of interest was run against various databases of choice. 25 significant protein sequences known to be involved in the crystalline cellulose biodegradation process in *R. thermocellum* were screened for known domains against SMART and Pfam database. These sequences included dockerin type I cellulosome proteins and cellulosome anchoring protein cohesion subunits with almost twenty sequences having catalytic subunits. Self comparison was employed and the steps shown in the flow diagram (Fig. 1) represent the workflow for prediction of domains and the results are shown in the Table. 1.

**Fig. 1.**
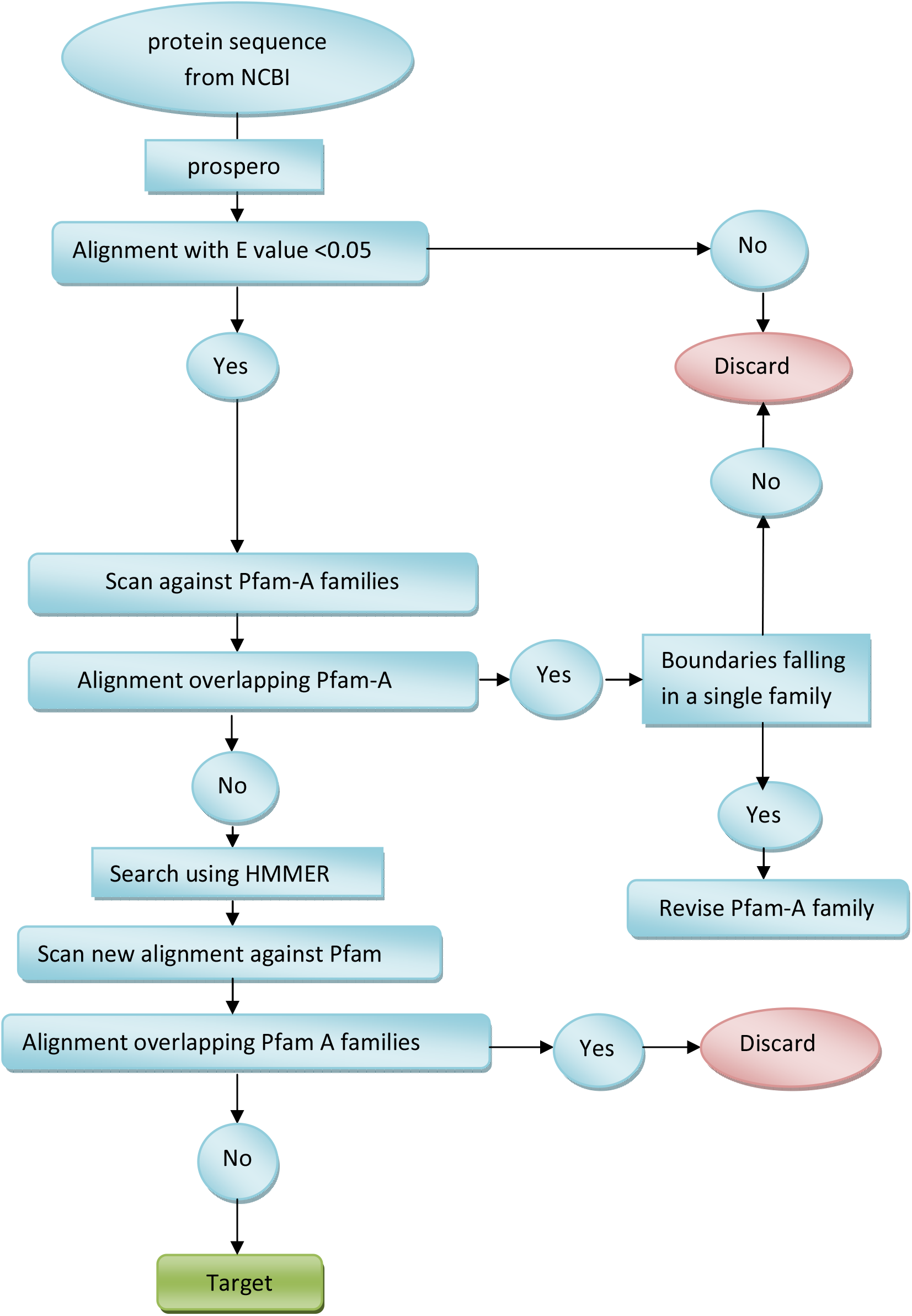
Domain hunt process to identify novel domains by detecting internal protein duplications.

**Table 1.** Novel domains identified in *R. thermocellum* cellulosome.

| Serial# | NCBI Accession# | Pfam-A detected domains | Insignificant Pfam-A detected domains | Pfam-B detected domains |
| --- | --- | --- | --- | --- |
| 1 | WP_003511666 | Dockerin_1 | - | - |
| 2 | WP_003511554 | Cohesin domain, S-layer homology domain | - | - |
| 3 | WP_003519375 | Cohesin domain, S-layer homology domain | Myco_arth_vir_N (Mycoplasma virulence signal region), Syndecan domain | PB003281, PB011093, PB000216 |
| 4 | WP_020458018 | Cohesin domain, S-layer homology domain | - | PB017474 |
| 5 | WP_020458017 | Cohesin, CBM_3 (cellulose binding) domain | Cna_B_2 (Cna protein B-type domain), Dockerin_1 (Dockerin type I repeat) | - |
| 6 | WP_020457956 | Dockerin_1 (Dockerin type I repeat) | Lipase_GDSL_2 GDSL-like Lipase/Acylhydrolase family(clan CL0264), DUF303 Domain of unknown function | - |
| 7 | WP_003518346 | Dockerin_1 (Dockerin type I repeat) | Gal-bind_lectin Galactoside-binding lectin Domain CL0004 clan | - |
| 8 | WP_003513507 | Dockerin_1 (Dockerin type I repeat) | TcdB_toxin_midN Insecticide toxin TcdB middle/N-terminal region CLO186 clan | - |
| 9 | WP_020457717 | CBM_6 (Carbohydrate binding module family 6), Dockerin_1 (Dockerin type I repeat) | - | - |
| 10 | WP_020457693 | Dockerin_1 (Dockerin type I repeat) | CBM_4_9 (Carbohydrate binding domain) | - |
| 11 | WP_020457693 | Dockerin_1 (Dockerin type I repeat), DUF285 Mycoplasma protein of unknown function | PKD (Polycystic Kidney Disease domain) | - |
| 12 | WP_011838264 | CarboxypepD_reg (Carboxypeptidase regulatory-like domain), Dockerin_1 (Dockerin type I repeat) | - | PB007732 |
| 13 | WP_003518268 | BNR (BNR/Asp-box repeat), Dockerin_1 (Dockerin type I repeat) | BNR_2 (BNR repeat-like domain), DUF4185 Domain of unknown function, Kelch_6 (Kelch motif) | PB000547 |
| 14 | WP_004463692 | Cohesin domain, SLH (S-layer homology domain) | DUF4398 Domain of unknown function | PB003281<br>PB005853 |
| 15 | WP_011837893 | Cohesin domain | DUF824 Salmonella repeat of unknown function | PB005853 |
| 16 | WP_003516262 | Cohesin domain | Spore_YhcN_YlaJ (Sporulation lipoprotein YhcN/YlaJ ) | - |
| 17 | WP_003516254 | Dockerin_1 (Dockerin type I repeat), DUF4353 Domain of unknown function | DUF4353 Domain of unknown function, Peptidase_C25_C (Peptidase family C25, C terminal ig-like domain) | - |
| 18 | WP_003516585 | Pectate_lyase_3 (Pectate lyase superfamily protein),dockerin_1 (dockerin type I repeat) | - | PB008307,<br>PB001086 |
| 19 | WP_011837833 | Cu_amine_oxidN1 (Copper amine oxidase N-terminal domain), Cohesin domain | - | - |
| 20 | WP_003519077 | Dockerin_1 (Dockerin type I repeat) | DUF843 Baculovirus protein of unknown function | - |
| 21 | WP_011837830 | Dockerin_1 (Dockerin type I repeat) | DUF1180 Protein of unknown function | PB001200 |
| 22 | WP_003512395 | Dockerin_1 (Dockerin type I repeat), RCC1 Regulator of chromosome condensation (RCC1) repeat | RCC1 (Regulator of chromosome condensation repeat ) | - |
| 23 | WP_003512367 | LTD (Lamin Tail Domain),<br>Fn3_assoc(Fn3 associated repeat),<br>CotH (CotH protein),<br>Dockerin_1 (Dockerin type I repeat domain) | - | PB013718 |
| 24 | WP_011837752 | Dockerin_1 (Dockerin type I repeat) | - | - |
| 25 | WP_003511967 | CotH (CotH protein family), Dockerin_1 (Dockerin type I repeat) | Integrin_b_cyt (Integrin beta cytoplasmic domain),<br>Metal_resist domain | - |

## MATERIALS AND METHODS

### Sequence data

A set of 25 enzyme sequences involved in cellulose degradation from *R. thermocellum* ATCC 27405 were taken from the NCBI database. The accession numbers of sequence data are provided in Table. 1.

### Methodology

A cogent mechanism in the evolution of novel proteins is interior duplication. A self comparison of proteins is a potent way of taking benefit of these internal duplications, rendering greater sensitivity as compared to all-against-all searching [7]. The approach illustrated in figure 1 for domain discovery has been employed previously with success [6, 7]. The steps described in the flow diagram (Figure 1) define the procedure that we have executed to identify novel domains by detecting internal protein duplications.

### Step 1

A set of 25 enzyme sequences involved in cellulose degradation from *R. thermocellum* were taken and Psi-Blast [8] was carried out. Low complexity sequence portion or repeat regions were masked using 'seg' [9]. Each protein sequence was compared against itself using Prospero [10]. Highest scoring self-self matches are returned. Matches have an E-value score determining the significance of each alignment

### Step 2

Highest scoring matches were kept for each sequence and a series of filters were applied to purge matches doubtful to be novel domains. All protein sequences corresponding merely to Pfam-A families were then discarded from the set and the rest were then aligned using T-Coffee [11].

Any sequence overlapping a Pfam-A family from this set was also removed except in case of sub sequences occurring within the boundaries of single Pfam-A family. This sort of occurrence is a sign that the family has more than one domain or repeat and requires refinement. An overlap is characterized as having residues occurring in the test alignment as well as the Pfam-A family alignment.

### Step 3

HMMER [12] software was used for creating profile-HMMs by means of an iterative search method utilizing the alignments generated by Prospero [6, 10]. The iterations were carried out until convergence, then they were realigned with T-Coffee and a single round of searching was done. Iterative search process was repeated if any new family members were identified.

Overlapping sequence portions in Prospero alignment were removed. Profile HMMs were built in both local (fs) and global (ls) mode. The profile-HMMs obtained as a result were scanned against the SWISS-PROT and TrEMBL databases [13]. An inclusion threshold of 0.05 was selected and an alignment of homologues discovered was built from HMMER package [12] using hmmalign program. This alignment was re-compared to the Pfam-A database for detection of any similarities to known families by HMM search. This step caused the removal of distant homologues of already described families [6].

### Step 4

In order to find out genomic milieu of the domain positions in the *R. thermocellum*, the genome was viewed using the Artemis [14] genome viewer.

## Results and Discussion

Domains predicted via sequence data represent evolutionary conserved sequences instead of distinct protein structures; but experience demonstrates that they frequently correspond to such structures as well. This discovery has prompted domains to be considered as the elementary units of protein evolution [15]. Potential novel domains in the cellulosome of *R. thermocellum* have been investigated by rapid automatic identification using Pfam-B [16] and from an initial set of proposed domain targets, 12 novel domains were identified and listed in the table (Table 1).

Pfam-A detections in the cellulosome sequences included Dockerin_1 domain, Cohesin, S-layer homology domain, DUF285, CBM_3, Carbohydrate binding module family 6, Carboxypeptidase regulatory-like domain, DUF4353, Copper amine oxidase N-terminal domain, pectate_lyase_3, Regulator of chromosome condensation (RCC1) repeat, CotH protein, Fn3 associated repeat and Lamin Tail domain. The Pfam-B detections are also shown in the table by their Pfam unique accession identifiers. Insignificant Pfam detections are those with scores below the threshold.

The primary purpose of this study was to identify novel protein domains in the cellulosome of *R. thermocellum*. To manually probe every distinct protein is massively time-consuming venture, and it is also not possible to add considerable annotation to several families built. Hence, automatic methods of family building and detailed annotation were carried out. In order to discover these domains in the cellulosome of *R. thermocellum*, it was only necessary to investigate the potential sequences, others were discarded. The principal reason for this was that no match was found to other proteins. This implies that comparative scans like this one will be useful only after sequencing of sufficient number of genomes of organisms exhibiting cellulose-degradation activity and carrying cellulytic enzymes.

While basing prospective work on studies like this, it ought to be noticed that the results are sets of hypotheses rather than actual descriptions. However the success of such approaches cannot be downgraded as well. An investigator studying HHE-containing protein in the past would have scanty information concerning it; now three candidates for the functional or active site residues are recognized and cation-binding function is assigned to it which can further be tested in the lab (reviewed by [6]). When one constituent of the family is described, information can be conveyed to its associated members. Deposition of the families into Pfam augments this and additional investigations employing Pfam will annotate these domains automatically, boosting the knowledge and comprehension of the remarkable organism *R. thermocellum*.

## References

1. Williams, T., J. Combs, B. Lynn and H. Strobel. 2006. Proteomic profile changes in membranes of ethanol-tolerant *Clostridium thermocellum*. Applied and Environmental Microbiology. 74:422–432.

2. Arnold, L. D., N. Michael, J. H. D. Wu. 2005. Cellulase, Clostridia, and Ethanol. Microbiol Mol Biol Rev. 2005. 69 (1): 124–154.

3. Weimer, P., and J. Zeikus. 1977. Fermentation of Cellulose and Cellobiose by *Clostridium thermocellum* in the Absence and Presence of Methanobacterium thermoautotrophicum. Applied and Environmental Microbiology. 33: 289–297.

4. Bateman, A., and E. Birney. 2000. Searching databases to find protein domain organization. Advances in Protein Chemistry. 54:137–157

5. Murzin, A.G. 1992. Structural Principles for the Propeller Assembly of Beta-Sheets – the Preference for 7-Fold Symmetry. Proteins-Structure Function and Genetics. 14:191–201

6. Yeats, C., S. Bentley, and Bateman. A. 2003. New knowledge from old: in silico discovery of novel protein domains in Streptomyces coelicolor. BMC Microbiol. 3:3–23

7. Ponting, C.P., R. Mott, P. Bork, and R. R. Copley. 2001. Novel protein domains and repeats in Drosophila melanogaster: Insights into structure, function, and evolution. Genome Research.11:1996–2008

8. Altschul, S.F., T.L. Madden., A. A. Schaffer, J.H. Zhang, Z. Zhang, W. Miller, and D. J. Lipman. 1997. Gapped BLAST and PSI-BLAST: a new generation of protein database search programs. Nucleic Acids Research. 25:3389–3402

9. Wootton, J.C., and S. Federhen. 1993. Statistics of Local Complexity in Amino-Acid-Sequences and Sequence Databases. Computers & Chemistry. 17:149–163

10. Mott, R. 2000. [http://www.well.ox.ac.uk/rmott/ARIADNE/prospero.shtml]

11. Notredame, C., D.G. Higgins, and J. Heringa. 2000. T-Coffee: A novel method for fast and accurate multiple sequence alignment. Journal of Molecular Biology. 302: 205–217

12. Eddy, S.R. 1998. Profile hidden Markov models. Bioinformatics. 14:755–76

13. Bairoch, A., and R. Apweiler. 2000. The SWISS-PROT protein sequence database and its supplement TrEMBL in 2000. Nucleic Acids Research. 28:45–48

14. Rutherford, K., J. Parkhill, J. Crook, T. Horsnell, P. Rice, M. A. Rajandream, and B. Barrell. 2000. Artemis: sequence visualization and annotation. Bioinformatics. 16:944–945

15. Ponting, C.P., and R. R. Russell. 2002. The natural history of protein domains. Annual Review of Biophysics and Biomolecular Structure. 31:45–71

16. Bateman, A., E. Birney, L. Cerruti, R. Durbin, L. Etwiller, S. R. Eddy, S. Griffiths-Jones, K. L. Howe, M. Marshall and E. L. L. Sonnhammer. 2002. The Pfam Protein Families Database. Nucleic Acids Research. 30:276–280

